# rhapsodist: a reproducible Snakemake workflow for BD Rhapsody single-cell RNA-seq data

**DOI:** 10.64898/2026.04.28.721304

**Authors:** Giulia Moro, Jiayi Wang, Ataman Sendoel, Izaskun Mallona

## Abstract

BD Rhapsody is a widely used single-cell RNA-seq (scRNA-seq) platform to profile gene expression. Based on barcoded beads, processing BD Rhapsody experiments is demanding because of the multiple barcode versions and layouts, which makes it difficult to locate barcodes by scanning reads positionally.

We introduce `rhapsodist`, a reproducible and scalable Snakemake workflow to process BD Rhapsody whole transcriptome analysis (WTA) data from raw FASTQ reads. As a result, `rhapsodist` generates count tables in a HDF5-backed `SingleCellExperiment` format. It supports all generations of BD Rhapsody beads, regardless of the barcode allowlist version and the presence of a variable-length diversity inset. `rhapsodist` orchestrates STARsolo, kallisto/bustools, salmon/alevin, and the official BD Rhapsody CWL pipeline, enabling direct cross-aligner comparison. Optionally, sample tag demultiplexing, read downsampling, tunable linker-mismatch tolerance, and per-sample reports are provided.

We showcase a remarkable consistency across aligner results, and highlight their differences in performance (e.g., quantification, clustering, or speed). On simulated data all four aligners recovered true barcodes with perfect precision and recall and equivalent UMI counts. On HeLa cells (a cell line; enhanced beads), both per-cell correlations across aligners and pseudobulk correlations were remarkably high on every comparison (Pearson r of 0.89–0.96 and 0.94–0.98 per cell and pseudobulk, respectively). On a mouse epidermis dataset with a more complex transcriptional profile (e.g., with celltypes; on legacy beads), cell type pseudobulk correlations showed a similar consistency (Pearson r 0.92–0.98), pointing to calling robustness. As expected, pseudoaligners (kallisto/bustools and salmon/alevin) were best performers speed-wise, and BD’s workflow recovered a higher number of barcodes and UMIs.

`rhapsodist` is available at https://github.com/imallona/rhapsodist under GPLv3, tried on Linux with conda, and includes unit and integration tests.

## Introduction

BD Rhapsody [Shum et al., 2019] is a widely used scRNA-seq platform to profile gene expression and other data modalities in single cells. Like Chromium and DROPseq, BD Rhapsody uses barcoded beads coated with DNA oligonucleotides [Gao et al., 2020, Salcher et al., 2024]. These oligonucleotides contain a tripartite cell barcode (CB), a unique molecular identifier (UMI), and a poly-dT oligo that captures mature mRNAs via their poly-A tail.

The original BD Rhapsody beads (v1, or legacy) carried a single type of oligonucleotides, the poly-dT. These were replaced by enhanced beads, which add a second type of molecule (named template switching oligonucleotides, or TSOs), and modify the cell barcode structure of dT oligonucleotides by prepending a variable-length diversity inset. Independently, we also observe that barcode allowlists for each part of the tripartite CB differ across bead versions, with at least two publicly available sets, with 96- and 384-sequence panels. TSO barcodes are not sequenced during most BD Rhapsody experiments, which profile WTA only.

Applying standard scRNA-seq pipelines to BD Rhapsody beads is hindered by the diversity of bead versions and by the presence of a variable-length diversity inset in dT cell barcodes (Figure S1) on enhanced beads. Aligners that locate barcodes by fixed position (alevin, kallisto, zUMIs [Parekh et al., 2018]) cannot handle the inset out of the box. The kallisto bus `BDWTA` mode targets v1 bead barcodes, which lack the diversity inset present in enhanced beads. This is not the case for STARsolo [Kaminow et al., 2021], which supports enhanced beads natively through its `CB_UMI_Complex` mode, by anchoring barcodes to the fixed parts of the barcode.

To our knowledge, existing options for enhanced BD Rhapsody data include the vendor’s CWL pipeline (so-called Seven Bridges), OpenPipelines (https://openpipelines.bio/), UniverSC Battenberg et al. [2022] and zUMIs [Parekh et al., 2018]. In contrast, `rhapsodist` adds (1) scalable parallel processing across bead versions, (2) cross-aligner comparison within one run, and (3) reproducible analysis-ready objects, BD-specific diagnostic reports, and pinned software environments.

## Design and implementation

### Required inputs

rhapsodist reads a YAML configuration file with three required stanzas: (i) a reference genome FASTA, gene annotation (GTF), and cDNA transcriptome FASTA files, and a working directory; (ii) workflow settings, including the aligner(s) to be run, the cell-filtering strategy, the number of threads (parallelization), a memory upper limit in MB, and optional parameters for downsampling and mismatch tolerance during barcode calling; (iii) a list of samples, each naming its paired CB+UMI (R1) and cDNA (R2) FASTQ files and its presumed bead version via two orthogonal keys: `allowlist` (whether the 96 or 384 panel is used) and `diversity_insets` (`yes` or `no`, Figure S1). Sample-tag demultiplexing is opt-in per sample (`use_sampletags`); the demultiplexing species (`human or mouse`) is only required when it is enabled. Indices may be supplied as local paths or URLs; in the latter case `rhapsodist` downloads them to the working directory. We provide example configuration files as Supplementary Material.

### Bead version detection and standardization

WTA cell barcodes are organized as three 9-nt CB segments (CB1, CB2, CB3) separated by fixed linkers, followed by an 8-nt UMI (Figure S1). The layout arises from two design components: whether the 96 or 384 size panels are used for each CB part (giving 96^3^ = 884,736 or 384^3^ = 56,623,104 possible cell barcodes), and from the presence or absence of the variable-length diversity inset prepended to CB1. Each enhanced bead has a mixture of different dT oligos with different insets (none, A, GT, or TCA) with the purpose of increasing library complexity. Hence, the inset shifts downstream CB and UMI start positions by 0–3 nt between reads, complicating positional slicing. Legacy (v1) beads use linkers ACTGGCCTGCGA and GGTAGCGGTGACA, the 96 panel allowlist, and no inset.

Before any read standardization or alignment, `rhapsodist` guesses the bead version by scanning the first 10,000 R1 reads. The detection function computes the fraction of reads matching legacy and enhanced fixed linkers, and classifies the sample accordingly. Given users also specify the bead version, a warning is triggered when the declared `allowlist` and `diversity_insets` disagree To facilitate debugging, linker mismatch rates are written to the descriptive report, so version mis-specification or unexpected error rates is reported as part of the standard QC, alongside barcode rank and UMI distribution plots.

To ease alignment, `rhapsodist` removes the diversity inset (if present), the fixed parts between the CB segments, and the sequence after the UMI using `cutadapt` v4.9 [Martin, 2011] and relying on anchoring at the fixed parts. The user-specified `cb_umi_max_errors` is passed to `cutadapt` as -e and controls the number of mismatches tolerated on the fixed linker parts of the R1 anchor. The default is 0 (exact linker match). To help pick a value, `rhapsodist’s` QC report scans the first 10,000 R1 reads and reports the fraction of reads within each Hamming distance of both the v1 and enhanced linker templates (Figure S2). We keep `cb_umi_max_errors = 0` by default.

### Memory-efficient handling of observed and valid cell barcodes

After standardizing the CB and UMIs, `rhapsodist` builds a sample-specific observed allowlist from the standardized CB+UMI FASTQ (Figure 1A). For each read, positions 1–9, 14–22, and 27–35 are extracted as candidate CB1, CB2, and CB3 sequences respectively. A read contributes to the allowlist only if all three segments match entries in the corresponding allowlist file as provided by the vendor (and included within the `rhapsodist` codebase); either the 96-sequence or 384-sequence panel depending on bead version). The full concatenated 27-nt barcode (CB1+CB2+CB3) fulfilling these criteria is added to an observed barcode set.

**Figure 1.**
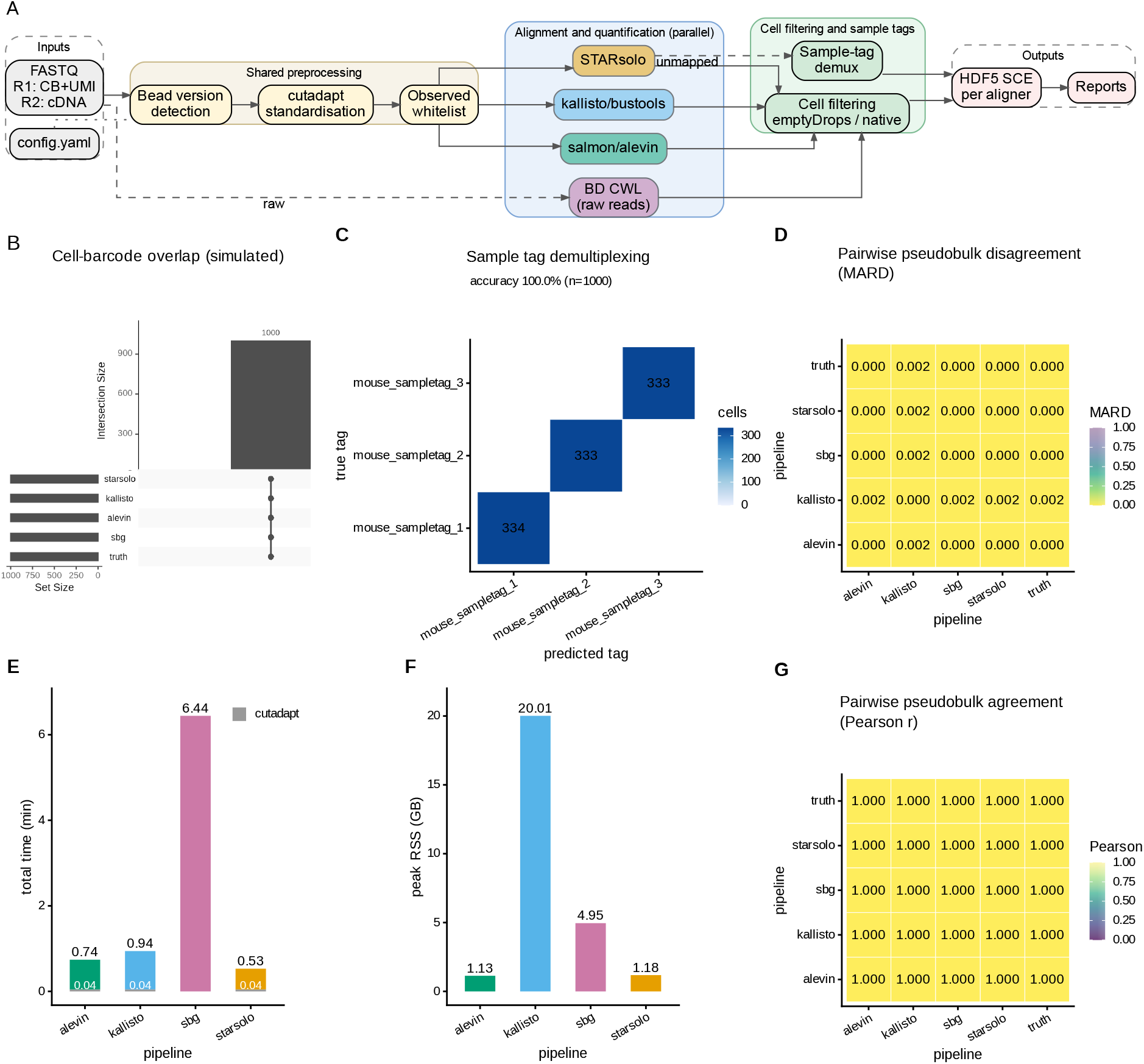
`rhapsodist` design and run on simulated BD Rhapsody data. (A) Workflow schematic: FASTQ reads pass through cutadapt barcode standardization and observed allowlist derivation before distribution to STARsolo, kallisto/bustools, salmon/alevin, and the BD Rhapsody CWL pipeline; each aligner emits an HDF5-backed SingleCellExperiment. (B) Cell barcodes recovered by each aligner against the simulated ground truth; all 1,000 simulated barcodes are recovered by every aligner. (C) Confusion matrix of predicted versus true sample tag across 1,000 simulated cells with three mouse tags; assignment accuracy is 100%. (D) Pairwise pseudobulk MARD (%) on simulated data, including ground-truth. (E) Total wall-clock time per aligner (minutes), excluding one-off `kallisto_install` and `bustools_install` rules and including the cutadapt timing (gray). (F) Peak resident memory per aligner (GiB). (G) Pairwise pseudobulk similary (Pearson r).

The resulting observed set (updated allowlist) replaces the full theoretical allowlist (up to 56 million entries for the 384 × 3 panel) with only those barcodes actually observed in the data. For real experiments this reduces the allowlist to tens of thousands of sequences, reducing memory usage and compute in subsequent steps.

### Downsampling

Experiments can be reproducibly downsampled by a downsample configuration parameter (a percentage of used reads in (0, 100], default 100). This is mainly useful for quick iterative runs on very deep libraries, or to make their library sizes comparable.

### Alignment and quantification

The SBG pipeline processes raw R1 reads, and other aligners use the standardized R1 reads and the observed whitelist. If the user selects more than one aligner, they run in parallel as independent tasks (Figure 1A).

`STARsolo` (STAR v2.7.10a [Dobin et al., 2013, Kaminow et al., 2021]) receives the standardized reads with fixed-position barcode geometry (`–soloType CB_UMI_Simple –soloCBstart 1 –soloCBlen 27–soloUMIstart 28 –soloUMIlen 8`). The genome index is built with `STAR –runMode genomeGenerate` using the user-supplied genome FASTA and GTF. UMI-deduplication strategy; multimapping strategy and cell filter are all configurable (defaults `Exact, Unique, EmptyDrops_CR`).

`kallisto v0.51.0/bustools` [Bray et al., 2016] are compiled from source at workflow runtime because their bioconda packages are not available on all platforms and architectures. kallisto maps the standardized reads against a transcriptome index and writes a BUS file; bustools sorts, corrects barcodes against the observed allowlist (up to 1 Hamming distance edit), and counts UMIs into a sparse matrix.

`salmon v1.10.3/alevin` [Srivastava et al., 2019]. Alevin receives the standardized reads with explicit geometry (`–bc-geometry ‘1[1-27]’ –umi-geometry ‘1[28-35]’ –read-geometry ‘2[1-end]’`) and the observed allowlist. A custom knee-point filter on alevin’s featureDump.txt separates real cells from empty droplets before the count matrix is loaded into R.

`BD Rhapsody CWL pipeline` (SevenBridges, v2.2.1 [Ulbrich et al., 2023, Li et al., 2024]; sbg). The official BD CWL workflow is run locally via cwl-runner on the original reads; a pre-built BD reference archive can be supplied or is built automatically from the STAR index and GTF. Unlike the other three aligners, `sbg` does not use the observed barcoded allowlist.

The `aligner` configuration key accepts any combination of the aligners above, encoded as `starsolo, kallisto, alevin`, and `sbg`, respectively.

### Cell filtering

rhapsodist supports two cell filtering strategies, configured via the `cell_filtering` field.

The `native` strategy uses each tool’s own filter. For STARsolo, this means the `soloCellFilter` mode set in the configuration (default `EmptyDrops_CR`, which applies the CellRanger 3.0 EmptyDrops variant). For alevin, the knee-point filter applied to `featureDump.txt` serves as the native filter. For kallisto/bustools, a knee-point filter from `DropletUtils::barcodeRanks` is applied to the bustools count matrix, falling back to the inflection point if the knee is undefined.

The `emptydrops` strategy applies DropletUtils `emptyDrops` [Lun et al., 2019] uniformly to all aligners. For STARsolo, the raw (unfiltered) count matrix is loaded and passed to `emptyDrops`. For kallisto, the bustools count matrix is used directly. For alevin, `emptyDrops` is applied to the knee-filtered barcode set, because alevin’s output omits most empty droplets and the knee filter further reduces the set before emptyDrops runs.

### Sample tag demultiplexing

Sample-tag demultiplexing is an opt-in. Enabled by the sample, it requires adding a `use_sampletags (yes or no`) and the demultiplexing `species (human or mouse`) panel, if enabled. A global key `skip_sampletags` disables the step for all samples.

When enabled, sample tags are antibody:oligo conjugates captured on dT barcodes. `rhapsodist` recovers sample tag reads from the reads unmapped by STARsolo: these reads failed alignment because their R2 sequence is not present in the genome. To further seed up sample tag quantification, only unmapped reads starting with the 25-nt common prefix shared by all sample tags (GTTGTCAAGATGCTACCGTTCAGAG) are analyzed. The variable region following the prefix is extracted and matched against the species-specific sample tag FASTA allowing some sequence variation. Each read is assigned to the closest tag, provided the Hamming distance on the variable region is at most two mismatches (default; configurable). Reads with no tag within this threshold are discarded. The output is a TSV with columns: cell barcode, UMI, assigned tag, and number of mismatches, and a demultiplexing report describing assignment rates and tags per cell distributions.

### Output and reports

Count tables are stored as HDF5-backed SingleCellExperiment (SCE) objects with metadata. HDF5 backing allows reading the count matrix while stored on disk rather than in memory, reducing memory overhead in commodity computers. The SCE objects follow Bioconductor guidelines and are compatible with downstream analysis tools such as scran, scater, and Seurat, and potentially imported by single-cell tools from the Python ecosystem.

Reports include: a *descriptive report* per aligner per sample showing barcode rank plots, genes and UMIs detected per cell, per-sample linker match rates against the declared chemistry (used as a QC panel for linker error rates or bead version mis-specification), and sample-tag counts (if applicable). If more than one aligner was used, a per-sample *cross-aligner comparison report* describes the number of cells, genes and UMIs per aligner, as well as pseudobulk agreement (MARD, Pearson r), and per-cell cross-aligner correlations; and a `performance report` describing, per run, the wall-clock time and peak memory (MiB) of each workflow step.

### Reproducibility and installation

`rhapsodist` requires conda (bioconda Grüning et al. [2018] and conda-forge) and Snakemake [Köster and Rahmann, 2012] and is written in R and python. R packages are installed into the conda environment at first run. kallisto and bustools are compiled from source at workflow runtime because their bioconda packages are not available on all platforms and hardware architectures.

The workflow can be installed as a Python CLI with `pip install –e`. and invoked as `rhapsodist –configfile configs/config.yaml –cores 10`, or plainly with `snakemake` commands directly using the `Snakefile`. Snakemake arguments such as `–rerun-incomplete` or `–nolock` can be appended directly to the `rhapsodist` CLI call. A simulation mode (`configs/sim_config.yaml`) generates and analyzes synthetic data providing a testing suit that completes in minutes. As for development, unit (pytest) and integration tests (GitHub actions) run on every commit.

### Simulation mode

A built-in synthetic BD Rhapsody WTA generator is provided to test the tool and benchmark it. By default, it produces a compact set of 1,000 cells, 100 genes, 50,000 empty droplets, and 100 sample-tag reads per cell across three mouse tags. Full parameters, the negative-binomial count model, and read simulation are detailed in Supplementary Methods and Figure S3.

### Application to experimental data

We showcase the application of rhapsodist to two experiments. First, to HeLa cells using enhanced v2 beads (384 whitelist, diversity inset present; SRA run `SRR35038871`), downsampled to 10% of paired reads [Moro et al., 2025]. Second, to a CROP-seq-based *in vivo* CRISPR screen originally designed to elucidate drivers of clonal expansion [Renz et al., 2024] in the mouse epidermis, using legacy v1 beads (96 whitelist, no inset; SRA `SRR24978231`, sample `sample_16_wta_p60`). Neither dataset carries sample tags, so `skip_sampletags` is set to `true`. The configurations used for both runs are described in the Supplementary Material. We disregard guide calling and perturbation-specific effects on this dataset.

HeLa and mouse reads were aligned to the GENCODE GRCh38 v46 and mouse GRCm39 vM36 references, respectively. Cell filtering used DropletUtils `emptyDrops` with FDR 0.001 [Lun et al., 2019] for the mouse skin sample and the tool-native knee-point filter for HeLa. Per aligner, a Seurat v5 object [Hao et al., 2024] was built, counts were log-normalized, PCA was computed on the 2,000 most variable genes, nearest neighbours were built on the first 30 principal components, clustering used the default Louvain algorithm with a 0.8 resolution, and the UMAP embedding was run with default parameters. For the skin sample, cell-type assignment used a marker-voting rule: each cell was assigned to the cell type with the highest sum of scaled expression across the markers described by the authors in their original paper Renz et al. [2024]. Cells below the threshold were left unassigned (`none` category). For HeLa, which is a clonal line with presumed homogeneity, we instead computed cell-cycle phase (G1, S, G2M) using Seurat to showcase cell-to-cell transcriptional variability. We quantified cross-aligner agreement with pairwise mean absolute relative difference (MARD) and Pearson correlation on pseudobulk log-normalized gene counts, per-cell Pearson r on log1p counts over shared barcodes and shared genes (downsampled to at most 500 cells per pair), and the adjusted Rand index (ARI) on Louvain partitions, cell-type labels (mouse skin), and cell-cycle phase labels (HeLa). Given Louvain cluster labels are arbitrary (cluster *i* in aligner A need not correspond to cluster *i* in aligner B), cluster-confusion matrices were reordered with the Hungarian algorithm using `clue::solve_LSAP` [Hornik, 2005].

Performance data were collected on a Linux server with an AMD EPYC 7742 processor.

## Results

We validated `rhapsodist` on three datasets. First, on a synthetic BD Rhapsody WTA dataset with known ground truth, used to verify barcode preprocessing, observed whitelist derivation, and per-aligner count recovery against an exact truth. Second, on HeLa cells using enhanced v2 beads (SRA `SRR35038871`), used to evaluate cell-to-cell quantification agreement on a clonal line where per-cell differences between aligners are not counfounded by cell types. And third, on a mouse epidermis dataset on legacy v1 beads [Renz et al., 2024] (SRA SRR24978231, used to explore cell filtering, marker-based cell typing, and cross-aligner agreement at the cluster and cell identity level.

### Validation on a synthetic BD Rhapsody dataset

The synthetic dataset contained 1,000 cells, 100 genes, 50,000 empty droplets, and 100 sample-tag reads per cell across three mouse tags (see Supplementary Methods). The simulation process generated the associated truths. The full simulation framework (parameters, count model, read synthesis, and outputs) is summarized in Figure S3. We ran STARsolo, kallisto/bustools, salmon/alevin, and the BD Rhapsody CWL pipeline (`sbg`).

For each aligner we asked four questions: does it recover the correct number of cells, does it detect every simulated gene, does it recover the exact set of true barcodes, and does it recover the simulated UMI counts per cell. All four aligners recovered 1,000 of 1,000 simulated barcodes with recall 1.00 and precision 1.00, and detected every one of the 100 simulated genes. Cell barcodes were identical across the four aligners and the simulated truth (Figure 1B). Median UMIs per cell matched the simulated value (153) for every aligner; the pseudobulk mean absolute relative difference versus truth was negligible, with 1.2 × 10^*−*5^ for sbg, 6.8 × 10^*−*5^ for STARsolo, 1.6 × 10^*−*4^ for salmon/alevin, and 2.3 × 10^*−*3^ for kallisto/bustools (Figure 1D).

Sample-tag demultiplexing on the simulated dataset assigned all 1,000 cells to their true tag (perfect accuracy, mean per-cell tag-read purity 1.00; Figure 1C).

Wall-clock time including 0.04 min for cutadapt was 0.83 min for STARsolo, 0.74 min for salmon/alevin, 0.94 min for kallisto/bustools, and 6.44 min for `sbg` (Figure 1E–F). Importantly, the `sbg` timings include the whole CWL flow and hence steps beyond alignment and barcode extraction, making a fair comparison difficult. One-off source-compilation rules (`kallisto_install` and `bustools_install`) were excluded from these totals because kallisto and bustools are built from source at first invocation and the resulting binaries are cached for later runs. Peak resident memory was 1.18 GiB (STARsolo), 1.13 GiB (salmon/alevin), 20.0 GiB (kallisto/bustools), and 4.95 GiB (`sbg`).

### Application to human HeLa (enhanced v2 beads)

We applied `rhapsodist` to a HeLa library on enhanced beads (384 whitelist, with diversity inset), and ran all four aligners against GENCODE human GRCh38 v46 with tool-native cell filtering. Because HeLa is a clonal line with reduced cell-to-cell variability, pairwise cell agreement is a proxy for alignment and quantification concordance. The four aligners called 4,357–5,453 cells with comparable UMIs, genes, and mitochondrial-fractions per cell (Figure 2A–B). Pseudobulk Pearson r ranged 0.94–0.98 on every aligner pair (Figure S4), while pseudobulk MARD spanned 18–74% (Figure 2C); the upper end corresponds to sbg pairs, and reflects the scale difference driven by its improved performance on quantifying UMIs). Across cells, per aligner Pearson r showed unimodal densities with median values 0.89–0.96 across the six aligner pairs (Figure 2D). As for comparing the similarity of predicted cell clusters, Louvain cluster partitioning was similar across aligners, with cluster ARIs ranging 0.63–0.80; and, in terms of predicted cell cycle phases (G1, S or G2M), their ARI ranged 0.69–0.77, pointing to consistency. Cell embeddings also depicted clusters and cell phases in similar relative positions (Figure 2E–H). Total end-to-end runtime was 12–89.7 min (including the cutadapt step when applicable), with a remarkable performance advantage for pseudoaligners, and peak resident memory 1.7–41.9 GiB (Figure 2I–J), with alevin being the top performer.

**Figure 2.**
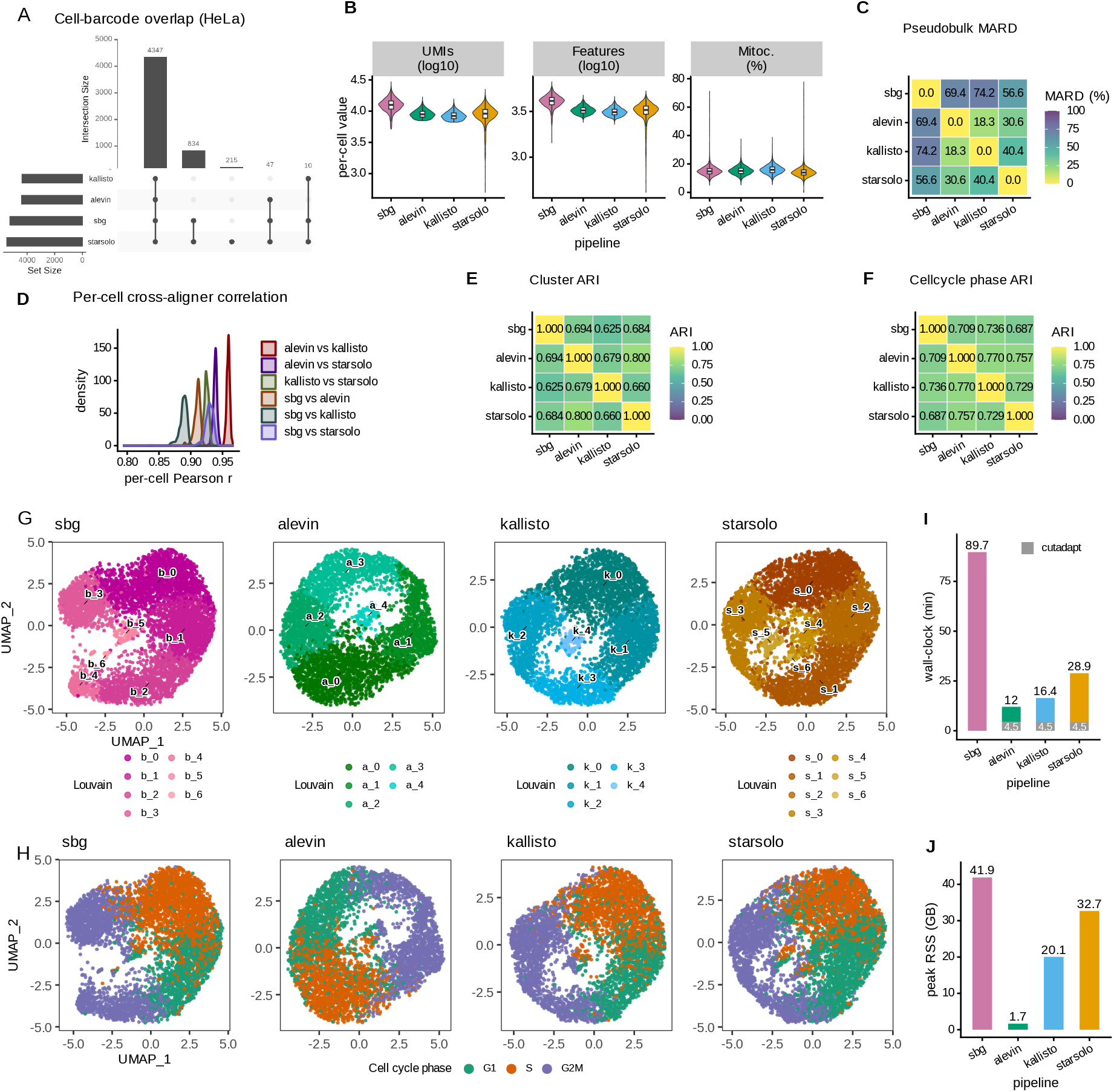
HeLa data using enhanced v2 beads (downsampled to 10% of the original). (A) Overlap on cell barcodes called by the four aligners after tool-native cell filtering. (B) Per-cell UMI count (log10), detected genes per cell (log10), and mitochondrial fraction (%). (C) Pairwise pseudobulk MARD (%) on log-normalized gene counts on shared cell barcodes. (D) Per-cell cross-aligner Pearson r on log1p counts over shared barcodes and shared genes, one density curve per aligner pair, up to 500 shared cells per pair. (E) Pairwise Louvain cluster ARI heatmap across aligners on shared barcodes. (F) Pairwise cell-cycle phase (G1, S, G2M) ARI heatmap across aligners. (G) UMAP embedding colored by Louvain cluster, one panel per aligner. (H) UMAP embedding colored by cell-cycle phase (as predicted by Seurat), one panel per aligner. (I) Total wall-clock time per aligner (minutes) and including the cutadapt timing (gray). (J) Peak resident memory per aligner (GiB). A pseudobulk Pearson r scatterplot is shown as Figure S4.

### Application to mouse epidermis (v1 beads)

We applied `rhapsodist` to sample `sample_16_wta_p60` from a postnatal mouse epidermis CRISPR screen [Renz et al., 2024] (v1 beads) with DropletUtils `emptyDrops` cell filtering (FDR 0.001) [Lun et al., 2019] against GENCODE mouse GRCm39 vM36. We neglected the effect of the guides and focused on transcriptional similarities by cell type. The four aligners called 40,546–54,982 cells with comparable per-cell UMI, gene, and mitochondrial-fraction distributions (Figure 3A–B), with consistenly higher values in detected UMIs and barcodes for `sbg`. Pseudobulk Pearson r on 33,812 shared gene symbols ranged 0.92–0.98 on every aligner pair (Figure S5). Pseudobulk MARD was 29–48% among alevin, kallisto and STARsolo and 89–92% against sbg, the high value again being driven by the higher number of UMIs read by SBG instead of by rank disagreement (Figure 3C). Pairwise Louvain cluster ARI was 0.81–0.86 and marker-based cell-type ARI 0.85–0.93 across all six aligner pairs (Figure 3E–H, confusion matrices in Figures S6 and S7). Coarser cell identity labels agreed more strongly than single-cell cluster labels: aligner choice shuffles clusters at the single-cell level but leaves downstream cell-type composition largely unchanged, matching prior reports for 10x Chromium data [You et al., 2021]. Total end-to-end runtime was 127.9–275.7 min across aligners (cutadapt step including, when applicable) and peak resident memory 6.5–34.6 GiB (Figure 3I–J).

**Figure 3.**
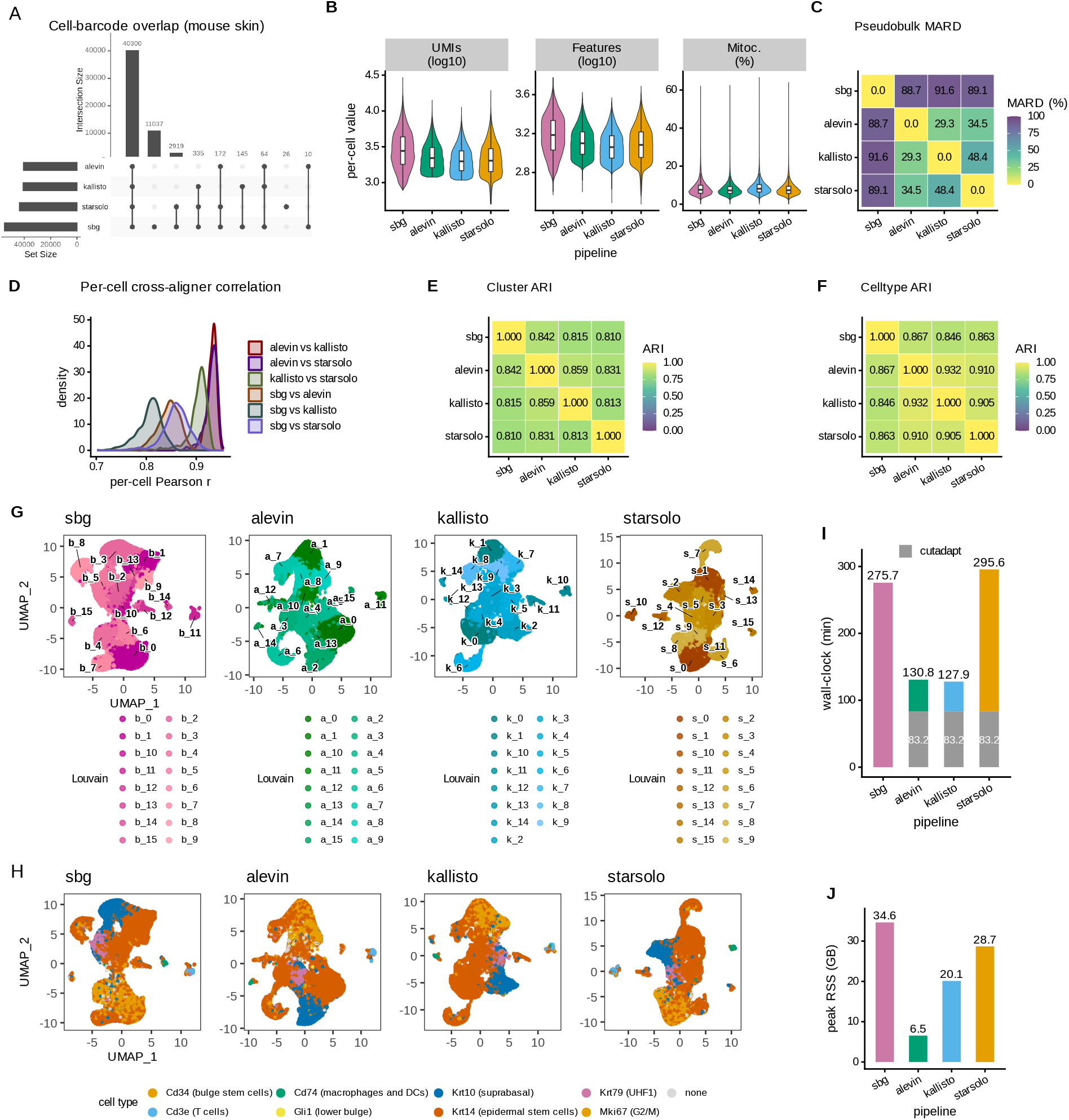
Mouse epidermis data (legacy beads). (A) Overlap of cell barcodes called by the four aligners after DropletUtils `emptyDrops`. (B) Per-cell UMI count (log10), detected genes per cell (log10), and mitochondrial fraction (%). (C) Pairwise pseudobulk MARD (%) on log-normalized gene counts on shared cell barcodes. (D) Per-cell cross-aligner Pearson r on log1p counts over shared barcodes and shared genes, one density curve per aligner pair, up to 500 shared cells per pair. (E) Pairwise Louvain cluster ARI heatmap across aligners on shared barcodes. (F) Pairwise marker-based cell-type ARI heatmap across aligners (confusion matrix shown as Figure S7). (G) UMAP embedding colored by Louvain cluster, one panel per aligner; confusion matrix shown at Figure S6. (H) UMAP cell embedding colored by marker-based cell-type assignment, one panel per aligner. (I) Total wall-clock time per aligner (minutes) including the cutadapt step (gray), where applicable. (J) Peak resident memory per aligner (GiB). A pseudobulk Pearson r scatterplot is shown as Figure S5.

### Availability and future directions

`rhapsodist` ships software environment files, a simulation run for testing, and documentation covering installation, configuration, and execution. Installation requires conda (bioconda and conda-forge channels) and Snakemake. The source code is available at https://github.com/imallona/rhapsodist under the GPLv3 terms; analysis scripts to build the figures in this paper are stored under `paper/`. The mouse epidermis and HeLa data are available at SRA `SRR24978231` [Renz et al., 2024] and `SRR35038871` [Moro et al., 2025], respectively. The GENCODE mouse GRCm39 vM36 and human GRCh38 v46 genome, annotation, and transcriptome releases are public and their download URLs are pinned in the `rhapsodist` configuration files. The synthetic dataset is generated by the run mode with configuration (`configs/sim_config.yaml`).

As main limitations, we only report two experimental datasets, one with legacy (v1) and another with enhanded beads, without sample tags. In terms of recovery of noise barcodes, we deliberately compare results from STARsolo, kallisto/bustools and salmon/alevin where cell barcodes and UMIs are recovered without any mismatch tolerance, whereas sbg runs handles and recovers mismatches internally. Allowing mismatches is built in on the software, and would raise per-cell UMI and gene counts, since 3% (HeLa) and 13% (mouse skin) of R1 reads match the linkers at exactly one mismatch (Figure S2). We expose `cb_umi_max_errors` as a user-tunable key and report the full Hamming-distance distribution per sample so the user can tune the parameter at will.

The current release has been tested only on Linux. Planned extensions include ATAC-seq co-processing, long read RNA-seq processing, and updates to incorporate future BD Rhapsody bead versions as their barcode layouts are released. Community contributions via pull request are welcome.

## Acknowledgements

We thank Anthony Sonrel for feedback on the workflow, and Mark D. Robinson for comments on the manuscript and access to compute.

## Author contributions

IM conceived the project and designed and wrote the workflow. IM, JW and GM analysed data. AS and GM provided data. IM wrote the manuscript and supervised the project. All authors edited and approved the final manuscript.

## Supplementary figures

**Figure S1:**
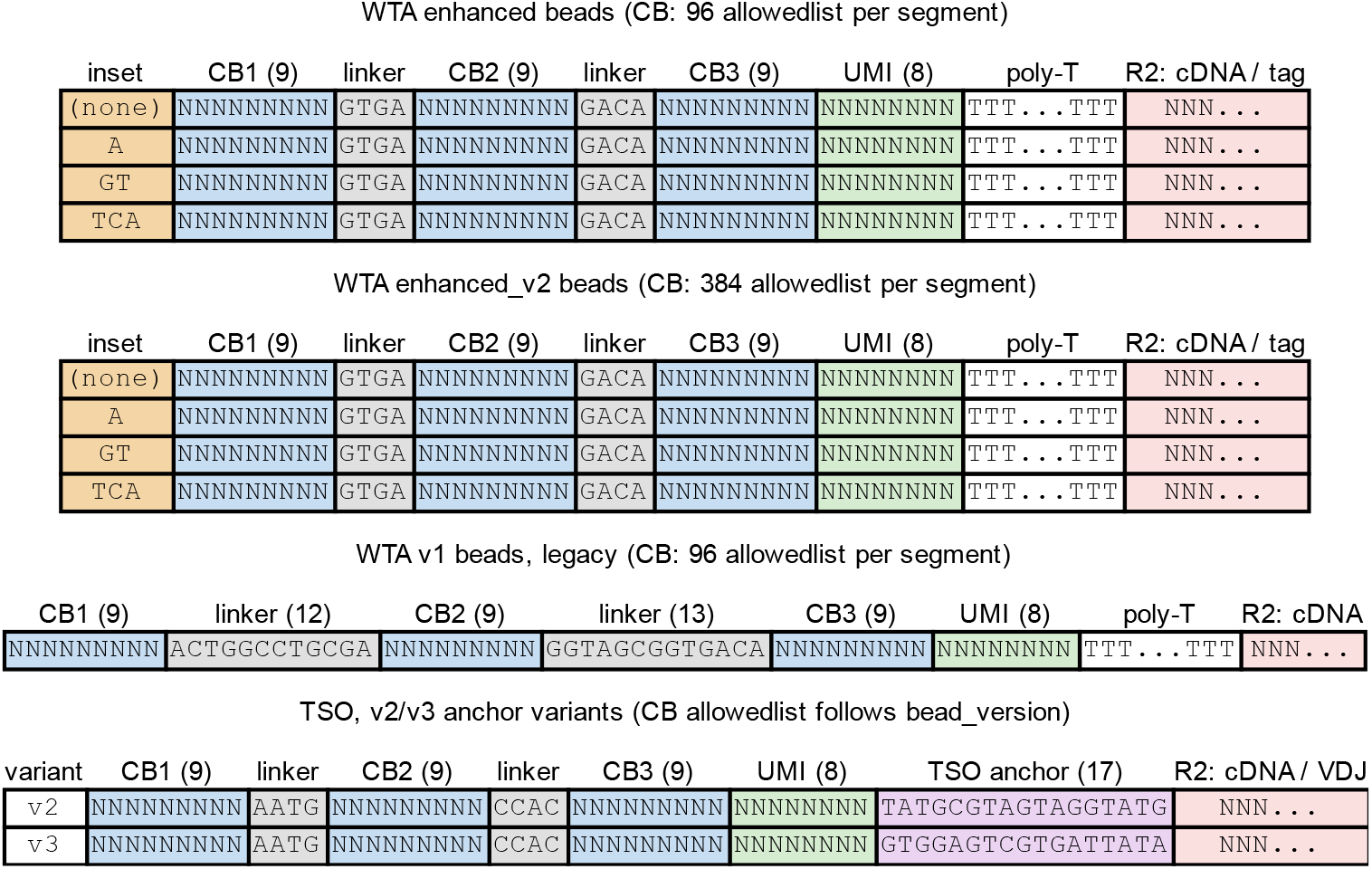
BD Rhapsody bead barcode structures handled by `rhapsodist`. Enhanced v2 dT barcodes, shown for each of the four diversity inset variants (`none, A, GT, TCA`). The three 9 nt CB segments (blue) are separated by the fixed linkers `GTGA` and `GACA;` an 8 nt UMI (green) follows CB3, then a poly-T stretch, then the cDNA or sample-tag sequence on R2. Legacy beads show fixed linkers ACTGGCCTGCGA and GGTAGCGGTGACA and no diversity inset. TSO barcodes share the tripartite CB+UMI layout but use linkers `AATG` and `CCAC` and carry no diversity inset; the TSO anchors differ in v2 (TATGCGTAGTAGGTATG) and v3 (GTGGAGTCGTGATTATA). TSO barcodes are not sequenced in most BD protocols. After sequencing, R1 reads the CB+UMI end; the minimum R1 length required to cover the full CB+UMI on enhanced beads is 3 + 9 + 4 + 9 + 4 + 9 + 8 = 46 nt. R2 reads the cDNA (or sample tag).

**Figure S2:**
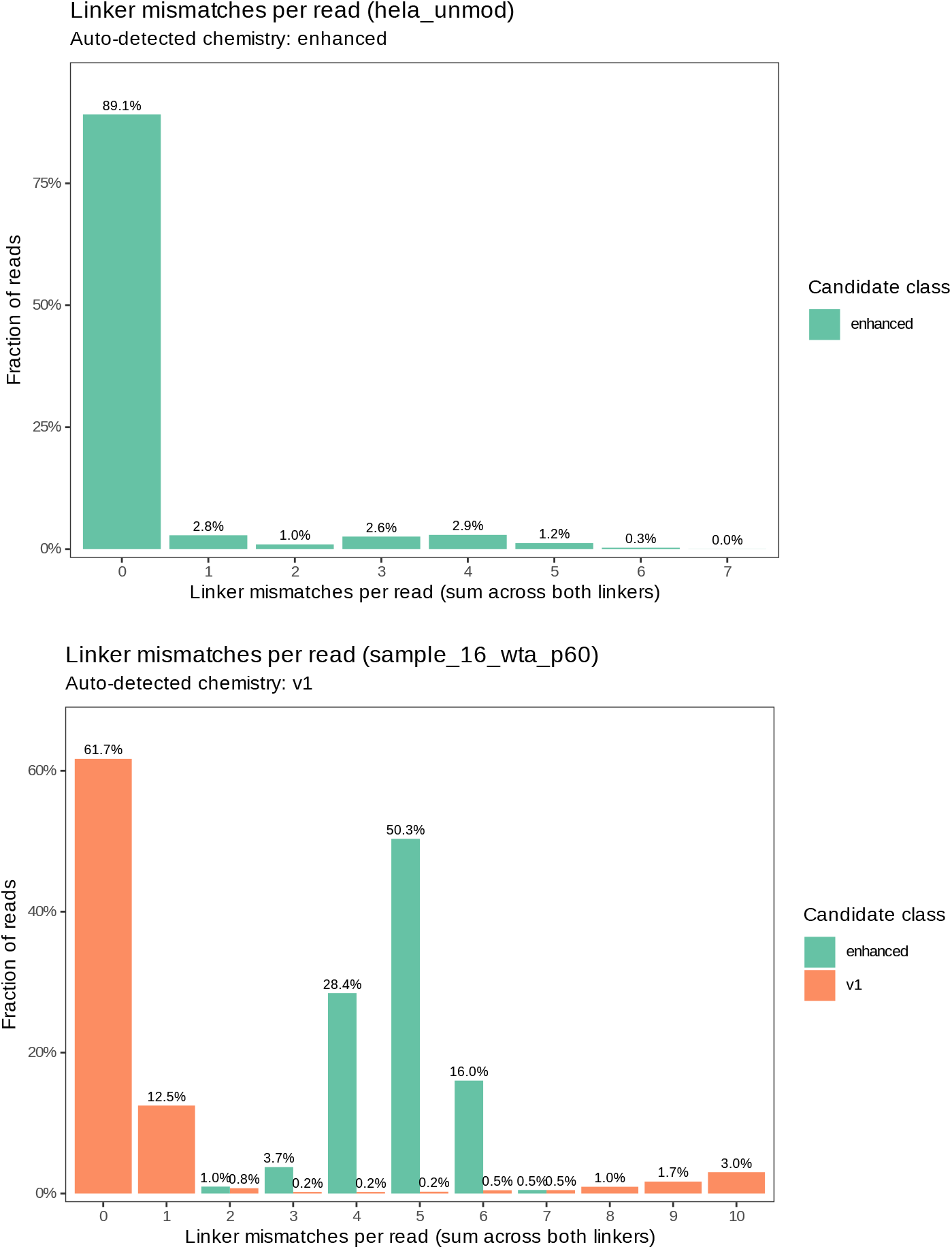
Per-read R1 linker-mismatch distributions on the first 10,000 reads of each real sample, scored against both v1 and enhanced linker templates. Bars: fraction of reads at each Hamming distance; color: candidate bead version. Top: HeLa (enhanced v2 beads): 89.1% of reads match the enhanced linkers exactly, 2.9% at one mismatch, 1.0% at two; the v1 template yields a broad non-zero distribution peaking near 19 mismatches (hence not plotted), consistent with auto-detection of the enhanced class. Bottom: mouse skin (v1 beads): 61.7% of reads match the v1 linkers exactly, 12.5% at one mismatch, less than 1% at two or three. The long tail at higher error counts on v1 is expected because the v1 fixed region spans 25 nt versus 8 nt for enhanced. The default setting running `rhapsodist` (`cb_umi_max_errors` value of 0, so exact linker match) retains the bulk of reads on both samples. Increasing it to 1 would potentially quantify an additional 2.9 % of HeLa and 12.5 % of mouse cells.

**Figure S3:**
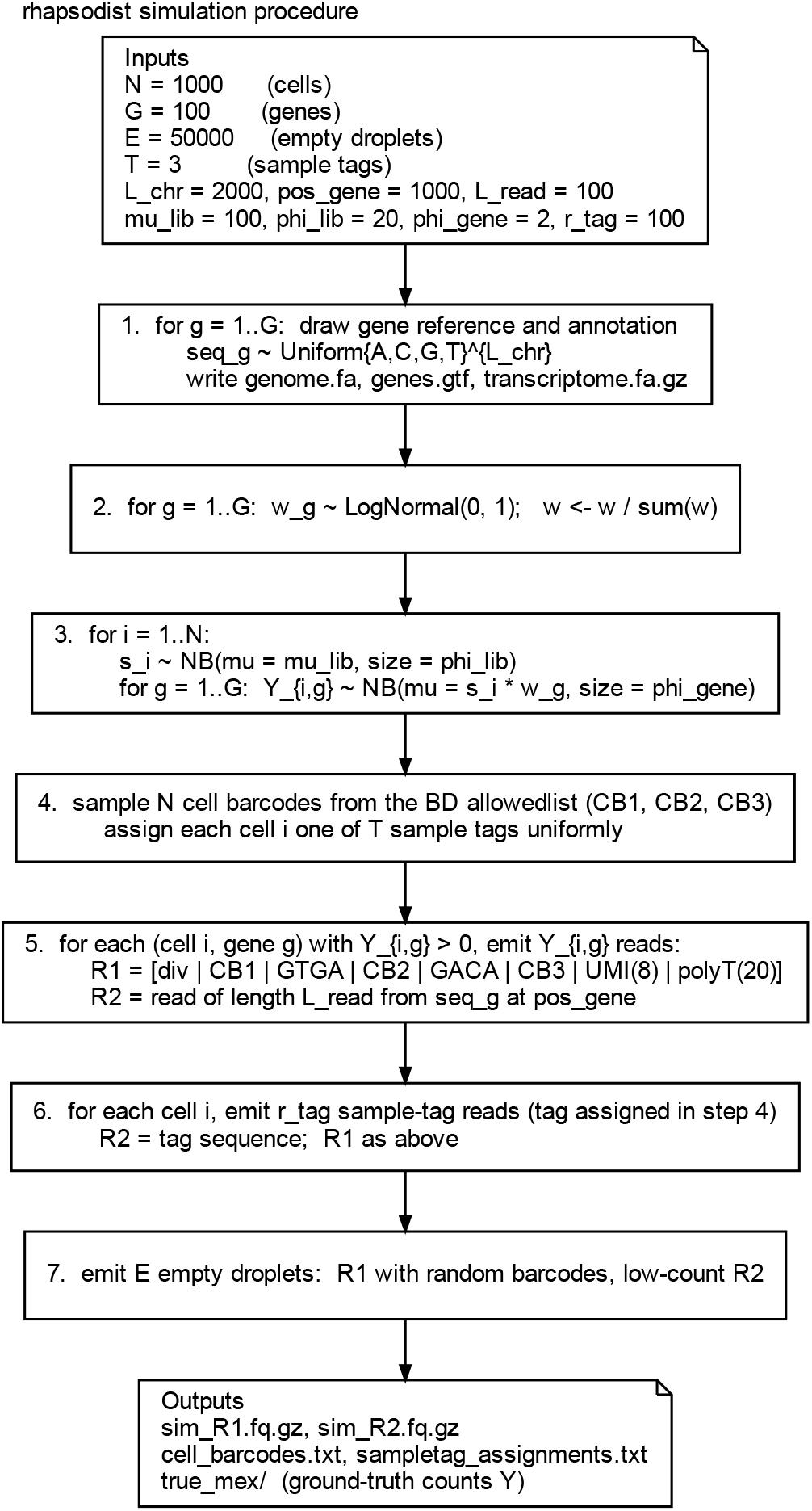
Summary of the `rhapsodist` simulation framework. Parameters (1,000 cells, 100 genes on 100 chromosomes, 50,000 empty droplets, three mouse sample tags, 100 sample-tag reads per cell) feed a negative-binomial count model: per-cell library sizes are drawn from NB(*µ*=100, size=20), per-gene weights are log-normal(0, 1) normalized to sum 1, and per-(cell, gene) counts are drawn from NB(*µ*=libsize×weight, size=2). Read synthesis writes a ground-truth matrix, cell barcodes, sample-tag assignments, and paired FASTQ reads (R1 encoding the diversity insert, CB1/CB2/CB3, UMI and polyT; and R2 carrying cDNA or sample-tag sequence).

**Figure S4:**
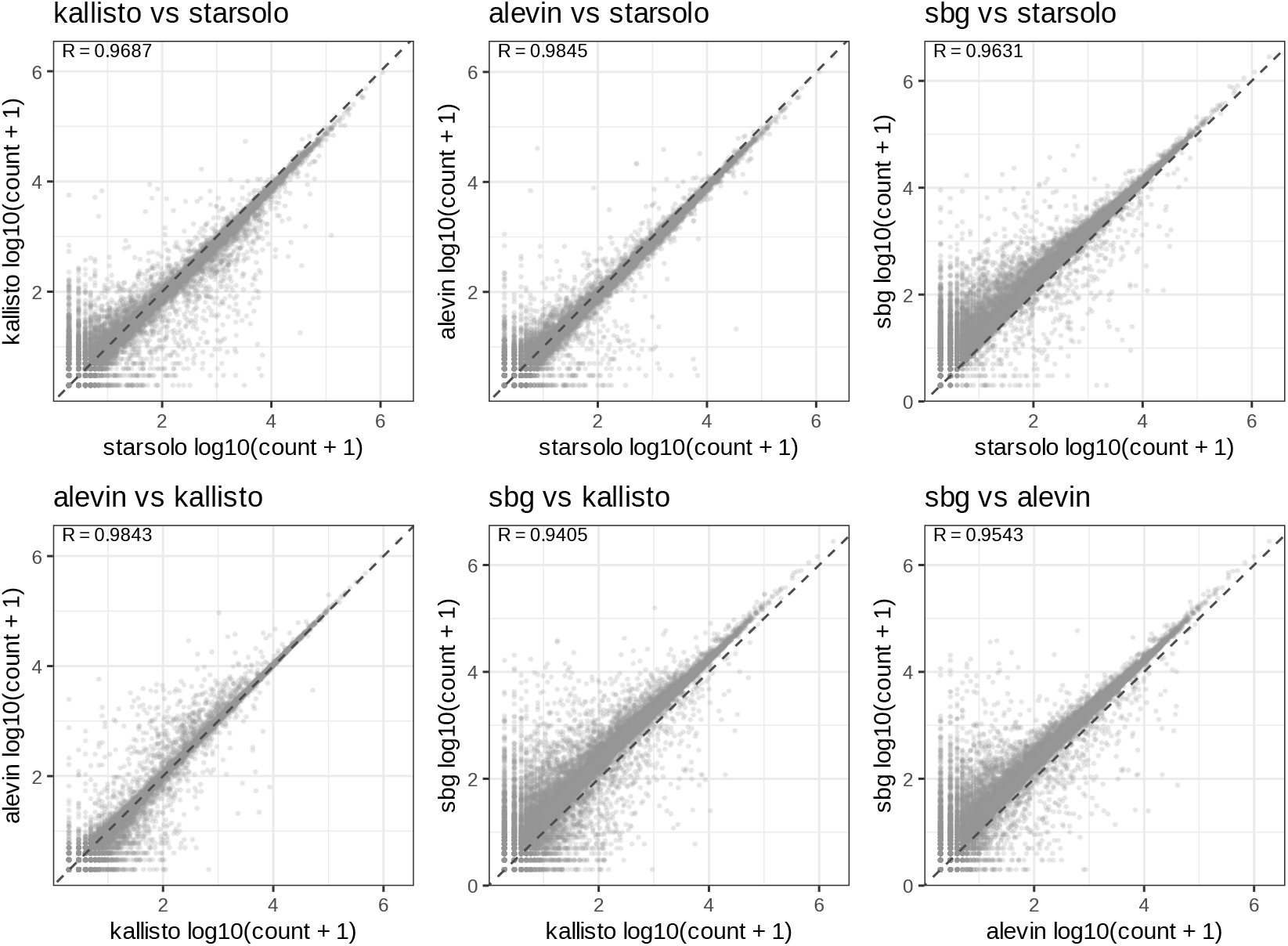
Pairwise pseudobulk scatterplots on HeLa data (enhanced v2 beads), across STARsolo, kallisto/bustools, salmon/alevin, and `sbg`. One panel per aligner pair; each point is a shared gene plotted as log10(count + 1); the dashed diagonal is y = x. Pearson r is annotated per panel.

**Figure S5:**
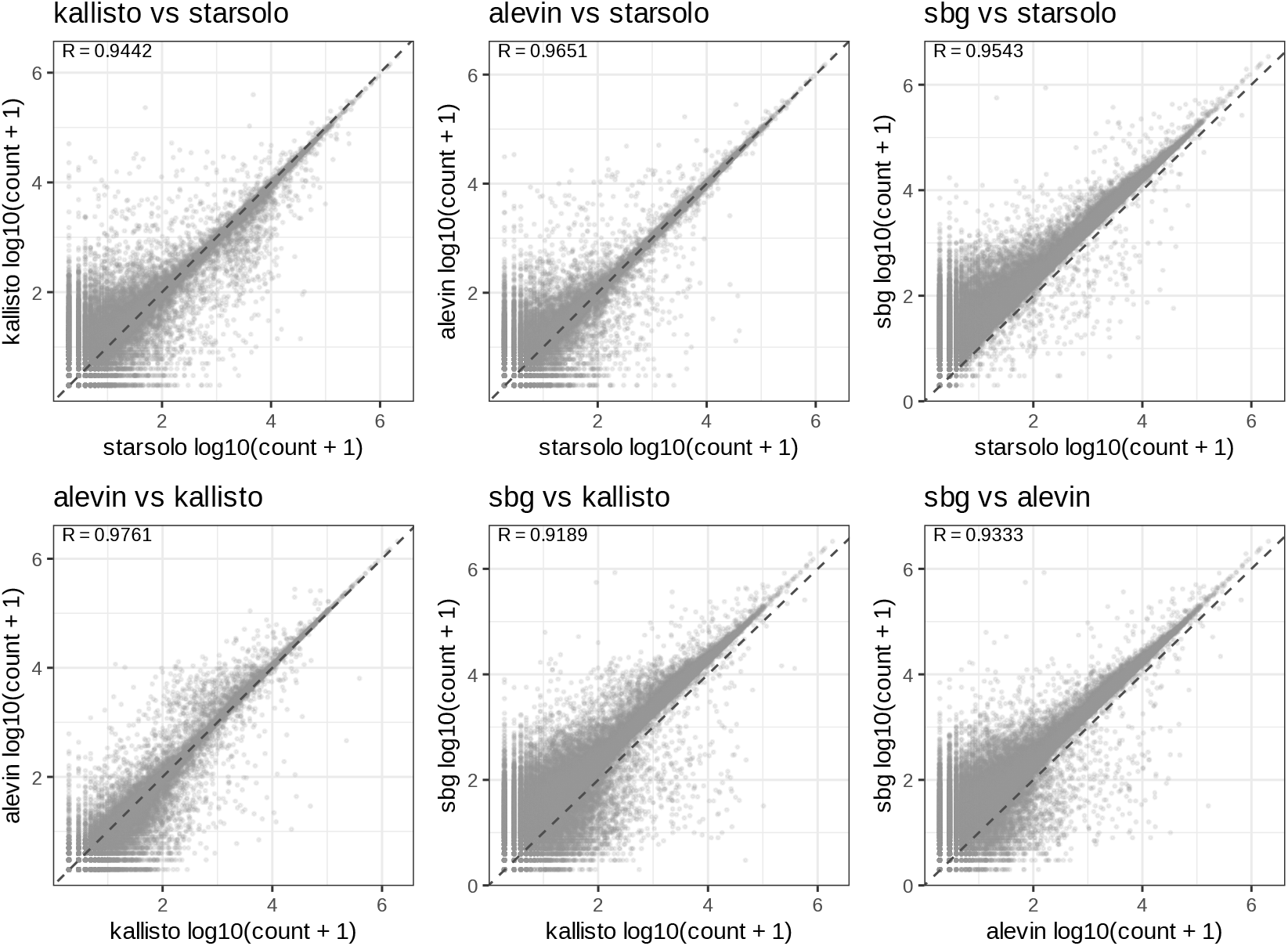
Pairwise pseudobulk scatterplots on mouse epidermis data (v1 beads), across STARsolo, kallisto/bustools, salmon/alevin, and `sbg`. One panel per aligner pair; each point is a shared gene plotted as log10(count + 1); the dashed diagonal is y = x. Pearson r is annotated per panel.

**Figure S6:**
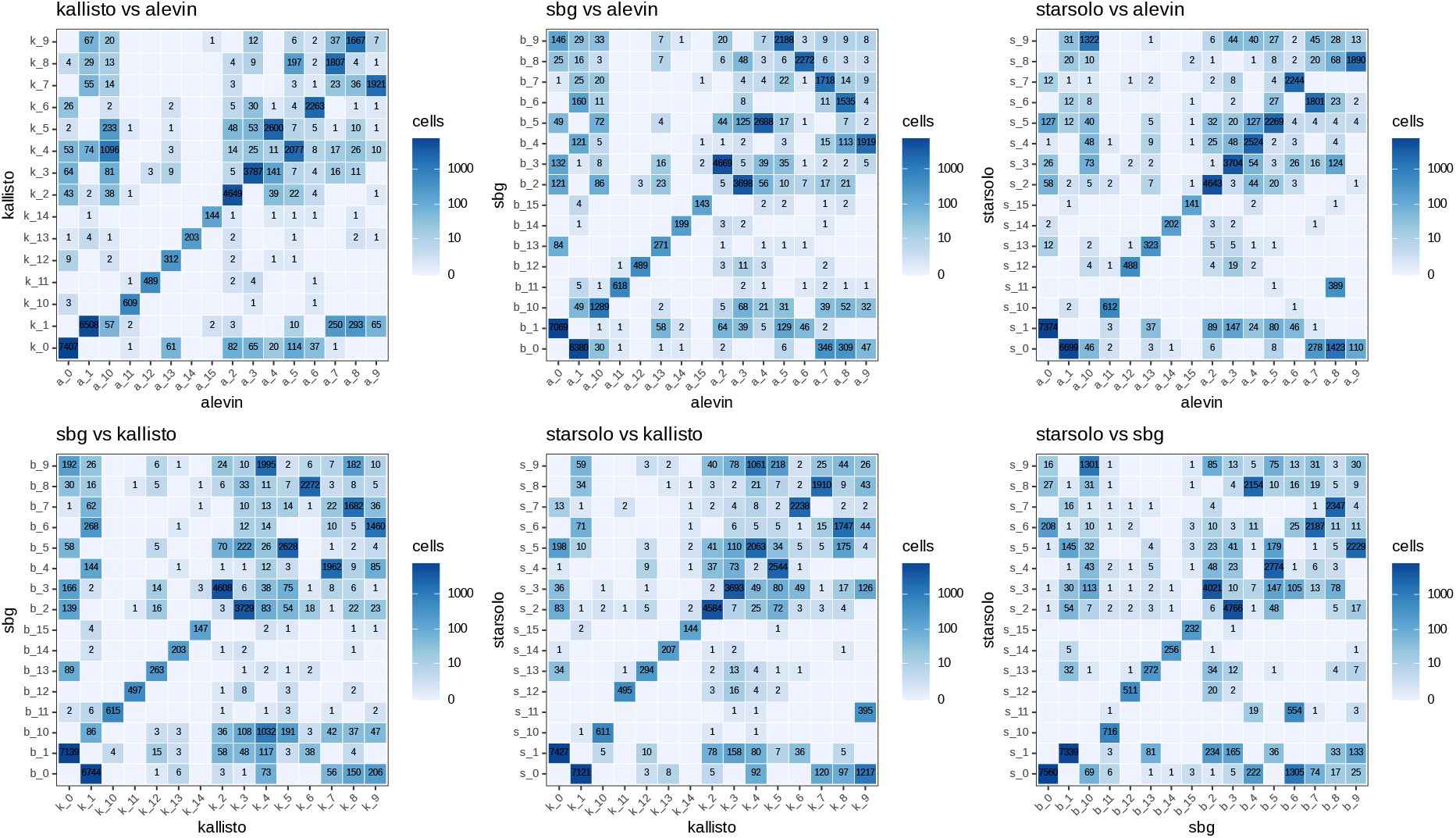
Pairwise confusion matrices for Louvain unsupervised clusters on the mouse skin dataset. Each heatmap cell shows the number of shared cell barcodes. Labels are reordered by Hungarian matching [Kuhn, 1955] on the pairwise cluster count table) so the dominant diagonal is visible. Reordering affects the displayed diagonal only, not the ARI values summarized in Figure 3E, which are invariant to label permutation.

**Figure S7:**
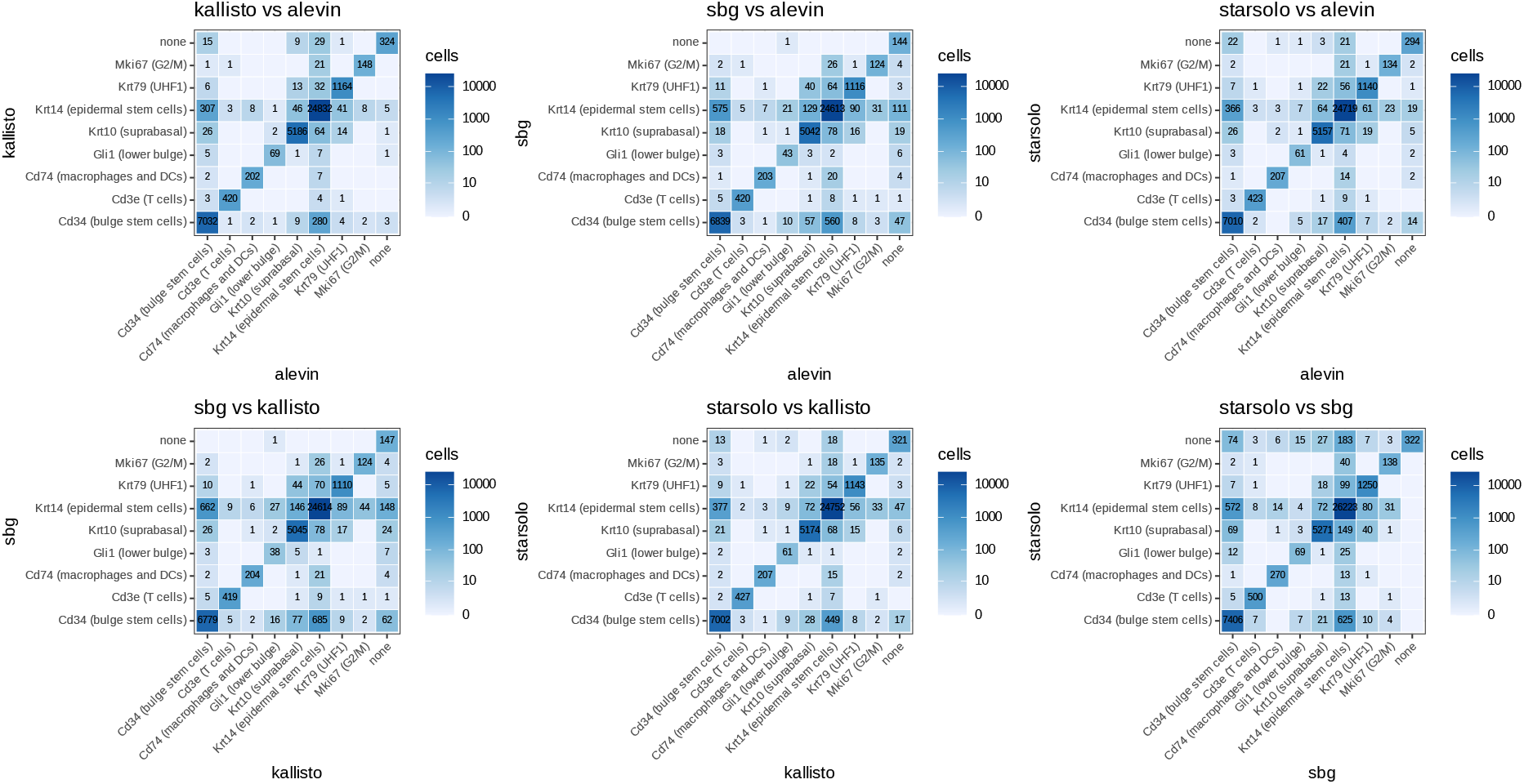
Pairwise marker-based cell-type confusion matrices on the mouse skin dataset. Each heatmap cell depicts the number of shared barcodes on log1p fill scale.

## Supplementary methods

The default simulation configuration (`configs/sim_config.yaml`) generates 1,000 cells, 100 genes, 50,000 empty droplets, and 100 sample-tag reads per cell across three mouse tags assigned in cyclic order (tags 1, 2, 3, …). The reference consists of 100 chromosomes of 2,000 bp each, with one unique 100 bp gene sequence embedded at position 1,000 on each chromosome, so mapping is unambiguous and counts can be verified exactly.

Per-cell library sizes are drawn from NB(*µ* = 100, size = 20), giving approximately 25% coefficient of variation across cells. Per-gene expression weights are drawn from LogNormal(0, 1) and normalized to sum to one. Final per-cell, per-gene counts are drawn from NB(*µ* = cell total × gene fraction, size = 2) restricted to a minimum of 1 per (cell, gene) pair so every simulated cell expresses every simulated gene; this restriction increases the per-cell median above the original *µ* of 100 (obseved median of 153). Each empty droplet receives between one and ten reads from a random gene to produce a bimodal barcode count distribution that is usable by alevin’s knee-point filter. Read length is 100 bp throughout; all base qualities are set to Phred 40 (I) to remove quality trimming as a source of variation during alignment/pseudoalignment. R1 carries the full enhanced v2 barcode structure (random diversity inset from {`none, A, GT, TCA`}, three 9 nt CB segments with `GTGA` and `GACA` linkers, 8 nt UMI, 20 nt poly-T tail); R2 contains the cDNA sequence. The random seed is fixed. The ground-truth count matrix is written in MEX format. The process is depicted in Figure S3.

### Example configuration files

Two annotated configurations are shipped with the workflow and cover both supported bead chemistries. `configs/sendoel2024_config.yaml` drives the mouse run on legacy v1 beads (96 whitelist, no diversity inset) and `configs/moro_mallona2025_config.yaml` drives the human HeLa run on enhanced v2 beads (384 whitelist, diversity inset present) with SBG enabled and 10% paired-read downsampling. Both fetch reads by SRA accession and download the GENCODE reference by URL, caching it under `reference_dir` for reuse across samples:

Listing 1: Mouse configuration (URL and SRA form).

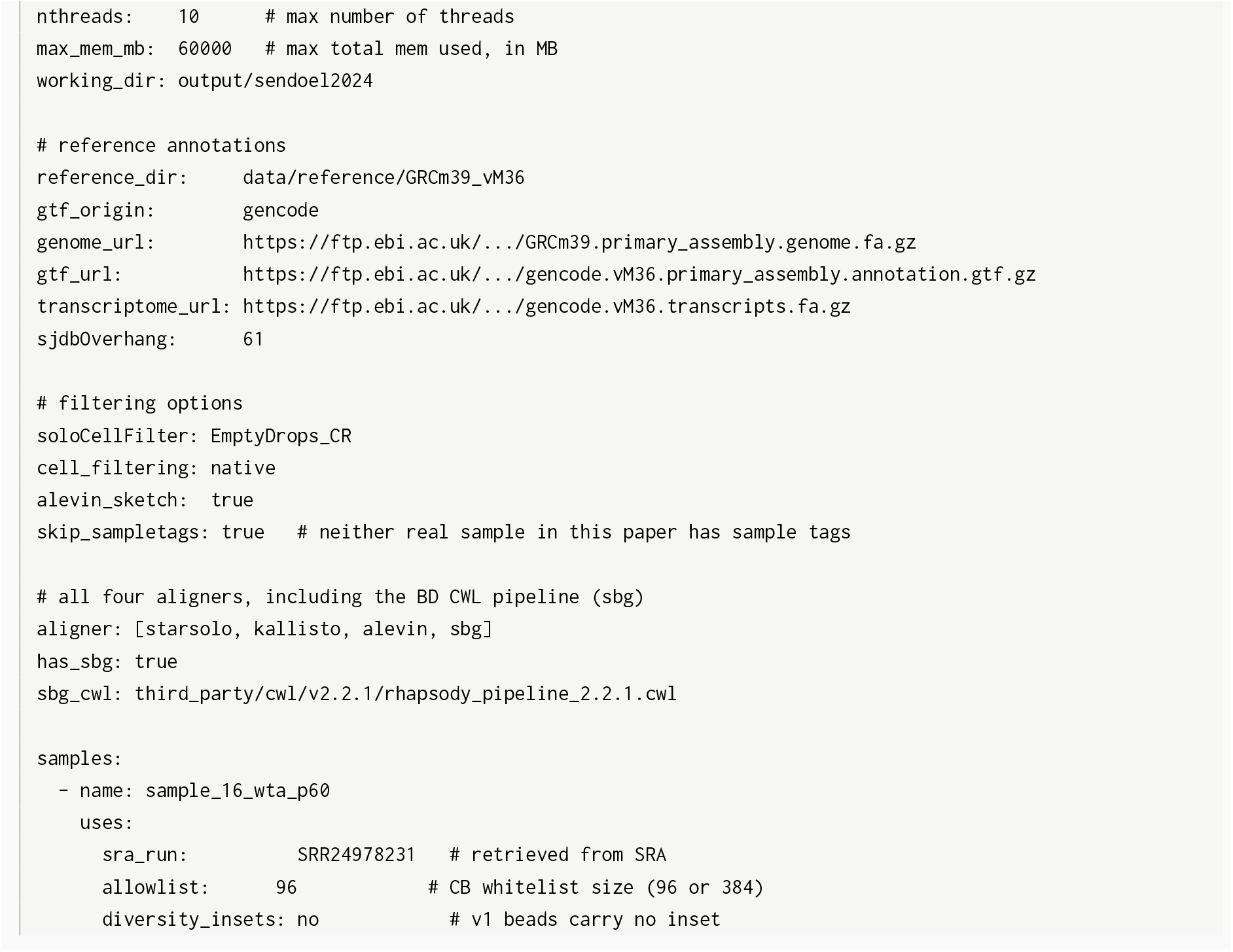

Listing 2: HeLa (enhanced v2 beads) configuration with SBG and 10% downsampling.

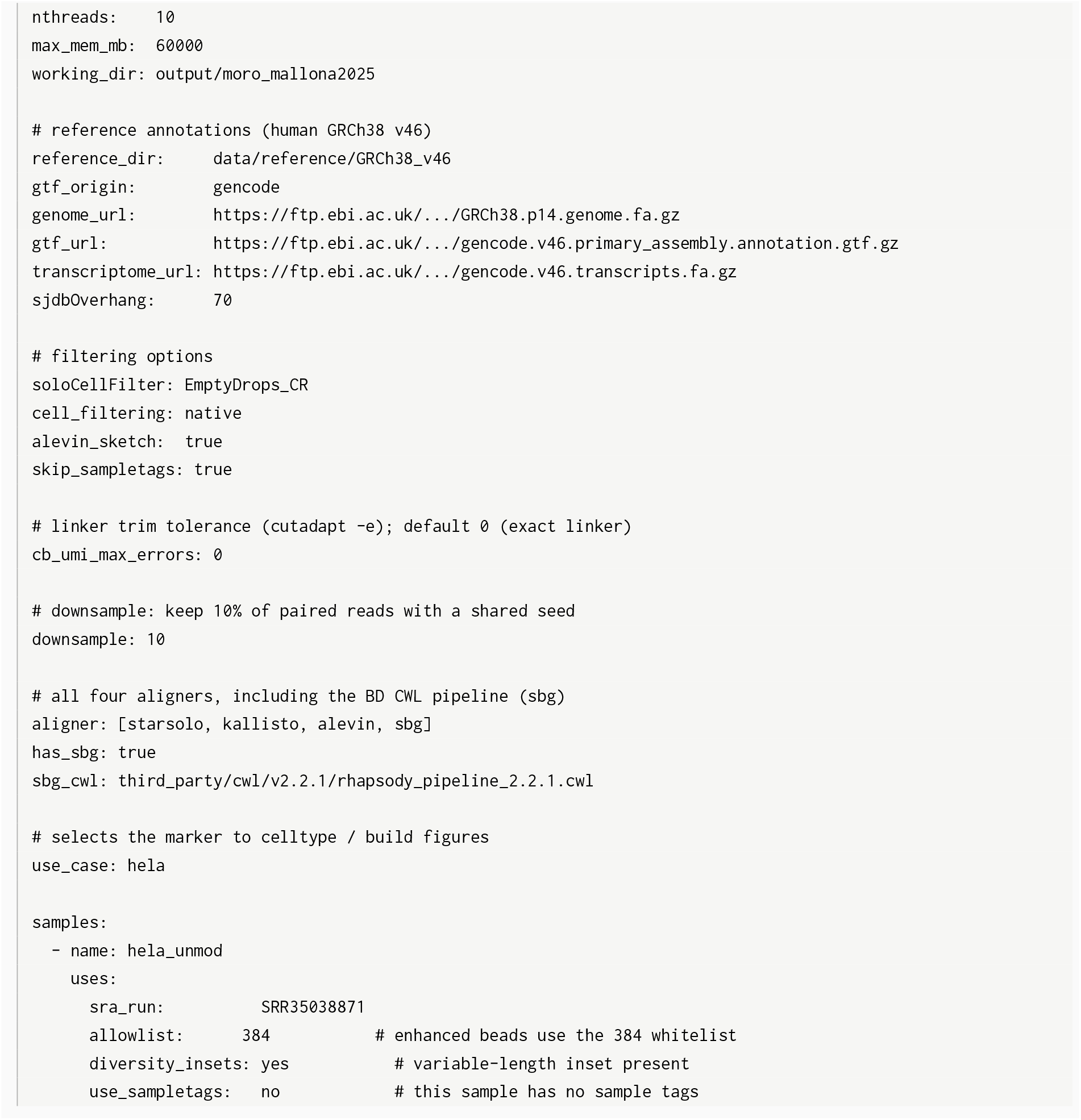

If the FASTQ files and the reference are already on disk, the same run can be expressed by replacing the `*_url` entries with local paths (`enome, gtf, transcriptome`) and replacing the per-sample `sra_run` for `cb_umi_fq` and `cdna_fq` pointing to the cell barcode and cDNA fastqs, respectively. The workflow detects local-path inputs automatically and skips the corresponding download rules:

Listing 3: Mouse configuration (local data variant).

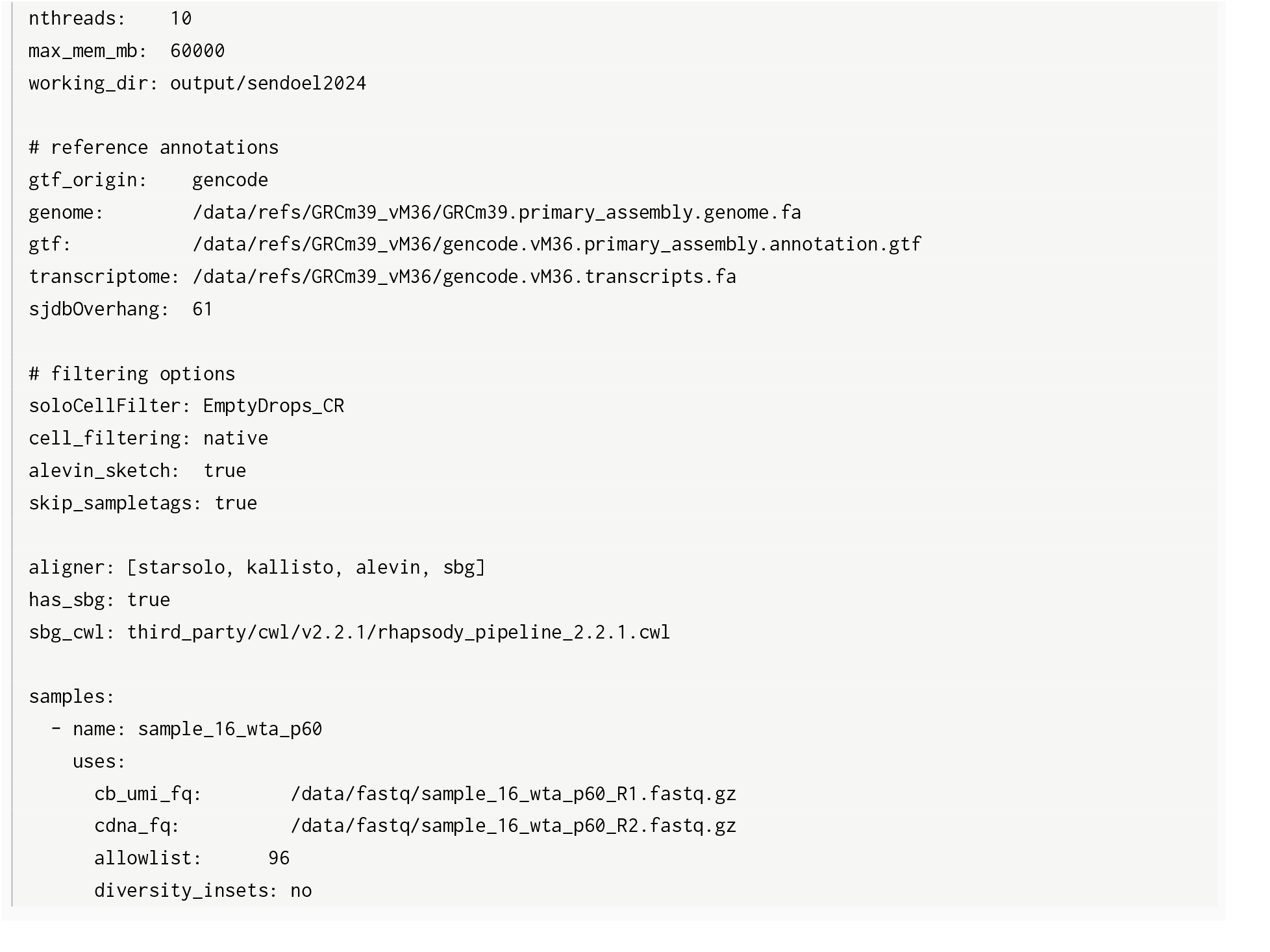

To run the workflow:

Listing 4: Running rhapsodist.

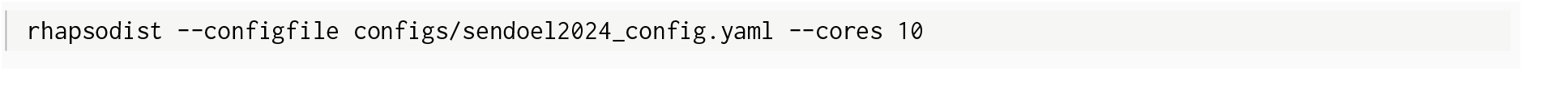

Outputs are written under `<working_dir>/<sample>/`, including HDF5 `SingleCellExperiment` objects per aligner, a cross-aligner comparison report, and the benchmark files used for the Results figures of this paper. Neither real sample in this paper carries sample tags, so `skip_sampletags` is set to `true;` the demultiplexing step is exercised instead by the built-in simulation (Figure 1C). For consistency, GTF and cDNA transcriptome must derive from the same GENCODE release and genome assembly.

## References

Kai Battenberg, S Thomas Kelly, Radu Abu Ras, Nicola A Hetherington, Makoto Hayashi, and Aki Minoda. A flexible cross-platform single-cell data processing pipeline. Nature Communications, 13(1): 6847, 2022.

Nicolas L Bray, Harold Pimentel, Páll Melsted, and Lior Pachter. Near-optimal probabilistic rna-seq quantification. Nature biotechnology, 34(5):525–527, 2016.

Alexander Dobin, Carrie A Davis, Felix Schlesinger, Jorg Drenkow, Chris Zaleski, Sonali Jha, Philippe Batut, Mark Chaisson, and Thomas R Gingeras. Star: ultrafast universal rna-seq aligner. Bioinformatics, 29(1):15–21, 2013.

Caixia Gao, Mingnan Zhang, and Lei Chen. The comparison of two single-cell sequencing platforms: Bd rhapsody and 10x genomics chromium. Current genomics, 21(8):602–609, 2020.

Björn Grüning, Ryan Dale, Andreas Sjödin, Brad A Chapman, Jillian Rowe, Christopher H Tomkins-Tinch, Renan Valieris, Johannes Köster, and Bioconda Team. Bioconda: sustainable and comprehensive software distribution for the life sciences. Nature methods, 15(7):475–476, 2018.

Yuhan Hao, Tim Stuart, Madeline H Kowalski, Saket Choudhary, Paul Hoffman, Austin Hartman, Avi Srivastava, Gesmira Molla, Shaista Madad, Carlos Fernandez-Granda, et al. Dictionary learning for integrative, multimodal and scalable single-cell analysis. Nature biotechnology, 42(2):293–304, 2024.

Kurt Hornik. A CLUE for CLUster Ensembles. Journal of Statistical Software, 14(12), 2005. doi: 10.18637/jss.v014.i12.

Benjamin Kaminow, Dinar Yunusov, and Alexander Dobin. Starsolo: accurate, fast and versatile mapping/quantification of single-cell and single-nucleus rna-seq data. Biorxiv, pages 2021–05, 2021.

Johannes Köster and Sven Rahmann. Snakemake—a scalable bioinformatics workflow engine. Bioinfor-matics, 28(19):2520–2522, 2012.

Harold W. Kuhn. The Hungarian method for the assignment problem. Naval Research Logistics Quarterly, 2(1–2):83–97, 1955. doi: 10.1002/nav.3800020109.

Wenyan Li, Sajad Razavi Bazaz, Chelsea Mayoh, and Robert Salomon. Analytical workflows for single-cell multiomic data using the bd rhapsody platform. Current Protocols, 4(2):e963, 2024.

Aaron T. L. Lun, Samantha Riesenfeld, Tallulah Andrews, The Phuong Dao, Tomas Gomes, and John C. Marioni. EmptyDrops: distinguishing cells from empty droplets in droplet-based single-cell RNA sequencing data. Genome Biology, 20(1):63, 2019. doi: 10.1186/s13059-019-1662-y.

Marcel Martin. Cutadapt removes adapter sequences from high-throughput sequencing reads. EMBnet. journal, 17(1):10–12, 2011.

Giulia Moro, Izaskun Mallona, Malwine J Barz, Joël Maillard, Michael David Brügger, Hassan Fazilaty, Quentin Szabo, Tomas Valenta, Kristina Handler, Fiona Kerlin, Lorenz Bastian, Claudia D Baldus, Andreas E Moor, Robert Zinzen, Mark D Robinson, Erich Brunner, and Konrad Basler. Rock and roi: single-cell transcriptomics with multiplexed enrichment of selected transcripts and region-specific sequencing. Nature Communications, 16(1):10991, 2025.

Swati Parekh, Christoph Ziegenhain, Beate Vieth, Wolfgang Enard, and Ines Hellmann. zUMIs - a fast and flexible pipeline to process RNA sequencing data with UMIs. GigaScience, 7(6), 2018. ISSN 2047-217X. doi: 10.1093/gigascience/giy059. URL https://academic.oup.com/gigascience/article/doi/10.1093/gigascience/giy059/5005022.

Peter F Renz, Umesh Ghoshdastider, Simona Baghai Sain, Fabiola Valdivia-Francia, Ameya Khandekar, Mark Ormiston, Martino Bernasconi, Clara Duré, Jonas A Kretz, Minkyoung Lee, Katie Hyams, Merima Forny, Marcel Pohly, Xenia Ficht, Stephanie J Ellis, Andreas E Moor, and Ataman Sendoel. In vivo single-cell CRISPR uncovers distinct TNF programmes in tumour evolution. Nature, 632(8024):419–428, August 2024.

Stefan Salcher, Isabel Heidegger, Gerold Untergasser, Georgios Fotakis, Alexandra Scheiber, Agnieszka Martowicz, Asma Noureen, Anne Krogsdam, Christoph Schatz, Georg Schäfer, Zlatko Trajanoski, Dominik Wolf, Sieghart Sopper, and Andreas Pircher. Comparative analysis of 10x chromium vs. bd rhapsody whole transcriptome single-cell sequencing technologies in complex human tissues. Heliyon, 10(7):e28358, 2024. ISSN 2405-8440. doi: 10.1016/j.heliyon.2024.e28358. URL https://www.sciencedirect.com/science/article/pii/S2405844024043895.

Eleen Y. Shum, Elisabeth M. Walczak, Christina Chang, and H. Christina Fan. Quantitation of mRNA Transcripts and Proteins Using the BD Rhapsody™ Single-Cell Analysis System, pages 63–79. Springer Singapore, Singapore, 2019. ISBN 978-981-13-6037-4. doi: 10.1007/978-981-13-6037-4_5. URL https://doi.org/10.1007/978-981-13-6037-4_5.

Avi Srivastava, Laraib Malik, Tom Sean Smith, Ian Sudbery, and Rob Patro. Alevin efficiently estimates accurate gene abundances from dscrna-seq data. Genome Biology, 20(1):65, 2019.

Jannes Ulbrich, Vadir Lopez-Salmeron, and Ian Gerrard. BD Rhapsody™ Single-Cell Analysis System Workflow: From Sample to Multimodal Single-Cell Sequencing Data, pages 29–56. Number 2584. Springer US, 2023. ISBN 978-1-0716-2755-6. doi: 10.1007/978-1-0716-2756-3_2. URL https://doi.org/10.1007/978-1-0716-2756-3_2.

Yue You, Luyi Tian, Shian Su, Xueyi Dong, Jafar S. Jabbari, Peter F. Hickey, and Matthew E. Ritchie. Benchmarking UMI-based single-cell RNA-seq preprocessing workflows. Genome Biology, 22:339, 2021. doi: 10.1186/s13059-021-02552-3.

